# Protection from viral rebound after therapeutic vaccination with an adjuvanted DNA vaccine is associated with SIV-specific polyfunctional CD8 T cells in the blood and mesenteric lymph nodes

**DOI:** 10.1101/2020.09.16.299511

**Authors:** Hillary C. Tunggal, Paul V. Munson, Megan A. O’Connor, Nika Hajari, Sandra E. Dross, Debra Bratt, James T. Fuller, Kenneth Bagley, Deborah Heydenburg Fuller

**Affiliations:** Department of Microbiology, University of Washington, Seattle, WA, USA; Washington National Primate Research Center, Seattle, WA, USA; Profectus Biosciences, Baltimore, MD, USA

## Abstract

A therapeutic vaccine that induces lasting control of HIV infection has the potential to eliminate the need for lifelong adherence to antiretroviral therapy (ART). This study investigated the efficacy of a therapeutic DNA vaccine delivered with a novel combination of adjuvants and immunomodulators to augment T cell immunity in the blood and gut-associated lymphoid tissue. In SIV-infected rhesus macaques, a DNA vaccine delivered by intradermal electroporation and expressing SIV Env, Gag, and Pol, and a combination of adjuvant plasmids expressing the catalytic A1 subunit of *E. coli* heat labile enterotoxin (LTA1), IL-12, IL-33, retinaldehyde dehydrogenase 2 and the immunomodulators soluble PD-1 and soluble CD80, significantly enhanced the breadth and magnitude of Gag-specific IFN-*γ* T cell responses when compared to controls that were mock vaccinated or received the same DNA vaccine delivered by Gene Gun with a single adjuvant, the *E. coli* heat labile enterotoxin, LT. Notably, the DNA vaccine and adjuvant combination protected 3/5 animals from viral rebound, compared to only 1/4 mock vaccinated animals and 1/5 animals that received the DNA vaccine and LT. The lower viral burden among controllers during analytical treatment interruption significantly correlated with higher polyfunctional CD8^+^ T-cells (CD8^+^ T cells expressing 3 or more effector functions) in both mesenteric lymph nodes and blood measured during ART and analytical treatment interruption. Interestingly, controllers also had lower viral loads during acute infection and ART suggesting that inherent host-viral interactions induced prior to ART initiation likely influenced the response to therapeutic vaccination. These data indicate that gut mucosal immune responses combined with effective ART may play a key role in containing residual virus post-ART and highlight the need for therapeutic vaccines and adjuvants that can restore functional quality of peripheral and mucosal T cell responses before and during ART.

**Author Summary:** HIV has caused significant human disease and mortality since its emergence in the 1980s. Furthermore, although antiretroviral therapy (ART) effectively reduces viral replication, stopping ART leads to increased viral loads and disease progression in most HIV-infected people. A therapeutic vaccine could enable HIV-infected people to control their infection without ART, but none of the vaccines that were tested in clinical trials so far have induced long-lasting control of virus replication. Here, we used the SIV rhesus macaque model to test a therapeutic vaccine consisting of DNA expressing SIV proteins and a novel combination of adjuvants to boost virus-specific immune responses. We found that our vaccine strategy significantly enhanced SIV-specific T cell responses when compared to controls and protected 3/5 animals from viral rebound. We determined that lower levels of virus replication post-ART were associated with enhanced T cell immunity in the gut-draining lymph nodes and blood. Our study highlights the critical role of T cell immunity for control of SIV and HIV replication and demonstrates that a successful therapeutic vaccine for HIV will need to elicit potent T cell responses in both the blood and gut-associated tissues.

## Introduction

ART greatly reduces HIV replication and restores CD4^+^ T cell counts, thus preventing progression to AIDS and prolonging the lifespan of HIV-infected people [1]. However, ART alone is unable to eliminate the latent viral reservoir, which necessitates strict lifelong adherence to a daily ART regimen [2]. For most individuals, ART interruption will lead to a resurgence in viral replication within weeks [3]. However, continuous usage of ART can be prohibitively expensive and may result in side effects that discourage compliance [4, 5]. Furthermore, ART cannot fully reverse the immune dysfunction induced by HIV, particularly in the gut mucosa, that drives chronic immune activation and disease pathogenesis [6, 7]. These drawbacks are why a vaccine or cure are still urgently needed to bring an end to the HIV pandemic.

To this end, many different cure strategies are in development, including therapeutic HIV vaccines designed to enhance virus-specific T cellular and humoral immune responses to achieve control of virus replication without ART. Numerous therapeutic HIV vaccines have been tested, both in the SIV/SHIV nonhuman primate (NHP) model and in human clinical trials, including protein subunit [8, 9], live-attenuated [10], dendritic cell [11], viral vectored [12, 13], and DNA vaccines [14–16]. Unfortunately, none of these approaches have resulted in durable control of viremia in human clinical trials [17].

While there are many barriers to an effective HIV therapeutic vaccine, we contend that the lack of success thus far may be partially attributed to insufficient vaccine immunogenicity in the gut and gut-associated lymphoid tissues (GALT). The gut is a major site of HIV and SIV replication [18–20], resulting in depletion and functional alteration of gut mucosal CD4^+^ T cells, and loss of antigen-presenting cells and innate lymphocytes [21]. These events contribute to structural damage of the gastrointestinal (GI) tract and systemic translocation of GI microbial products that drive chronic immune activation and disease pathogenesis [22, 23]. Here, we investigated a strategy to enhance mucosal-associated immunity by incorporating adjuvants and immunomodulators specifically designed to potently boost HIV-specific immunity in both the periphery and the gut mucosa, with the goal of eliciting robust control of viremia following ART cessation.

We previously showed in the SIV rhesus macaque model that epidermal co-delivery of plasmids encoding an SIV whole antigen DNA vaccine and the mucosal adjuvant, heat-labile *E. coli* enterotoxin (LT), induced durable protection from viral rebound and disease progression for 10 months after ART withdrawal in 5/7 animals, in comparison to 1/7 animals in the mock vaccinated control group [15]. This outcome was associated with significant increases in SIV-specific CD8^+^ T cells expressing dual effector functions in the blood, and IFN-*γ* T cell responses in both the blood and gut [15]. Notably, mucosal T cell responses in vaccinated animals significantly correlated with reduced virus production in both mucosal tissues and in plasma [15], indicating that SIV-specific responses in the gut may be important for controlling viral rebound. These results demonstrate that inducing vigorous immune responses in the mucosa may be critical for control of viremia and the importance of using potent adjuvants to maximize vaccine efficacy.

These findings prompted us to explore methods to further enhance the breadth, function and magnitude of mucosal T cell responses. Using the SIV rhesus macaque model, we tested the therapeutic efficacy of a novel combination of adjuvants and a multiantigen SIV DNA vaccine (MAG) delivered by intradermal electroporation. Our adjuvant combination (AC) consisted of co-delivered plasmids encoding the catalytic subunit of LT (LTA1), the cytokines IL-12 and IL-33, the enzyme retinaldehyde dehydrogenase 2 (RALDH2), soluble PD-1 (sPD-1), and soluble CD80 (sCD80). LTA1 is a potent adjuvant that performs similarly to LT, through the recruitment and activation of dendritic cells [24, 25]. The IL-12 adjuvant has been widely used in both NHP and human clinical trials [26, 27] and promotes differentiation of naïve CD4^+^ and CD8^+^ T cells to Th1 and cytotoxic T lymphocytes (CTLs), respectively. IL-33 has also been shown to augment vaccine-induced immunity in mice [28, 29], and works by directly promoting the activity of Th1 cells and CTLs [30, 31]. RALDH2 was included to enhance vaccine immunogenicity in the mucosa through the conversion of retinaldehyde to retinoic acid, the molecule responsible for inducing the expression of the mucosal homing factors CCR9 and *α*4*β*7 on activated lymphocytes [32, 33]. Finally, since previous studies showed that blocking the PD-1 and CTLA-4 pathways can enhance antigen-specific immunity, reduce immune activation, and restore immune exhaustion [34, 35], we co-delivered plasmids expressing rhesus-specific sPD-1 and sCD80 to block the interaction of CD8^+^ T cells expressing PD-1 and CTLA-4 with antigen presenting cells (APCs) expressing PDL-1 and CD80.

The results shown here demonstrate that vaccination of SIV-infected, ART-treated rhesus macaques with the MAG DNA vaccine and the adjuvant combination (MAG + AC) increased the magnitude, breadth, and durability of IFN-*γ* T cell responses when compared to the mock vaccinated controls or animals vaccinated with the MAG DNA adjuvanted with only LT (MAG + LT), with the majority of the response targeting Gag sequences. Following analytic treatment interruption (ATI, discontinuation of ART), three out of five animals (60%) in the MAG + AC group were protected from viral rebound compared to only 20% in the MAG + LT group and 25% in the mock vaccinated group. A comparison of immune responses in animals that controlled viral rebound (controllers) to those that exhibited immediate viral rebound during ATI (noncontrollers), regardless of vaccine group, showed that controllers had higher polyfunctional SIV-specific CD8^+^ T cells in the mesenteric lymph nodes (MLN) and blood. Additional comparisons of controllers and noncontrollers showed the relative response to ART also correlated with viral control during ATI. Together, these findings highlight an important role for effective ART and mucosal and systemic CD8^+^ T cell responses in controlling viral rebound during ATI. These studies also suggest that inherent host responses to the virus that occur at the earliest stages of infection may determine the ability of an individual to respond to antiretroviral drug therapy and therapeutic vaccination.

## Results

### NHP Study Design

We previously showed that an SIV therapeutic DNA vaccine adjuvanted with LT induced protection from viral rebound in >70% of SIV-infected macaques and was associated with higher peripheral polyfunctional SIV-specific CD8^+^ T cells and broader specificity in the mucosal T cell response [15]. Here, we sought to bolster SIV-specific CD8^+^ T cell immunity in the gut-associated lymphoid tissues (GALT) through adjuvants demonstrated to enhance T cell immunity and mucosal homing [26, 29, 33], to further test the hypothesis that increasing mucosal immunity will improve efficacy of therapeutic vaccination.

Rhesus macaques were intravenously infected with the highly pathogenic, primary isolate SIVΔB670 [36]. At six weeks post-infection (wpi), animals began ART, consisting of emtricitabine (FTC), tenofovir (PMPA), and Raltegravir, administered daily, and starting at 32 wpi, macaques received a series of 5 DNA immunizations, spaced 4 weeks apart (Fig 1). Prior to initiating therapeutic immunizations, we stratified the animals so that each group had comparable levels of plasma viremia and blood CD4^+^ T cell counts (S1 Fig) during acute infection and ART to balance the effects of pre-existing virological and host factors among all groups.

**Fig 1.**
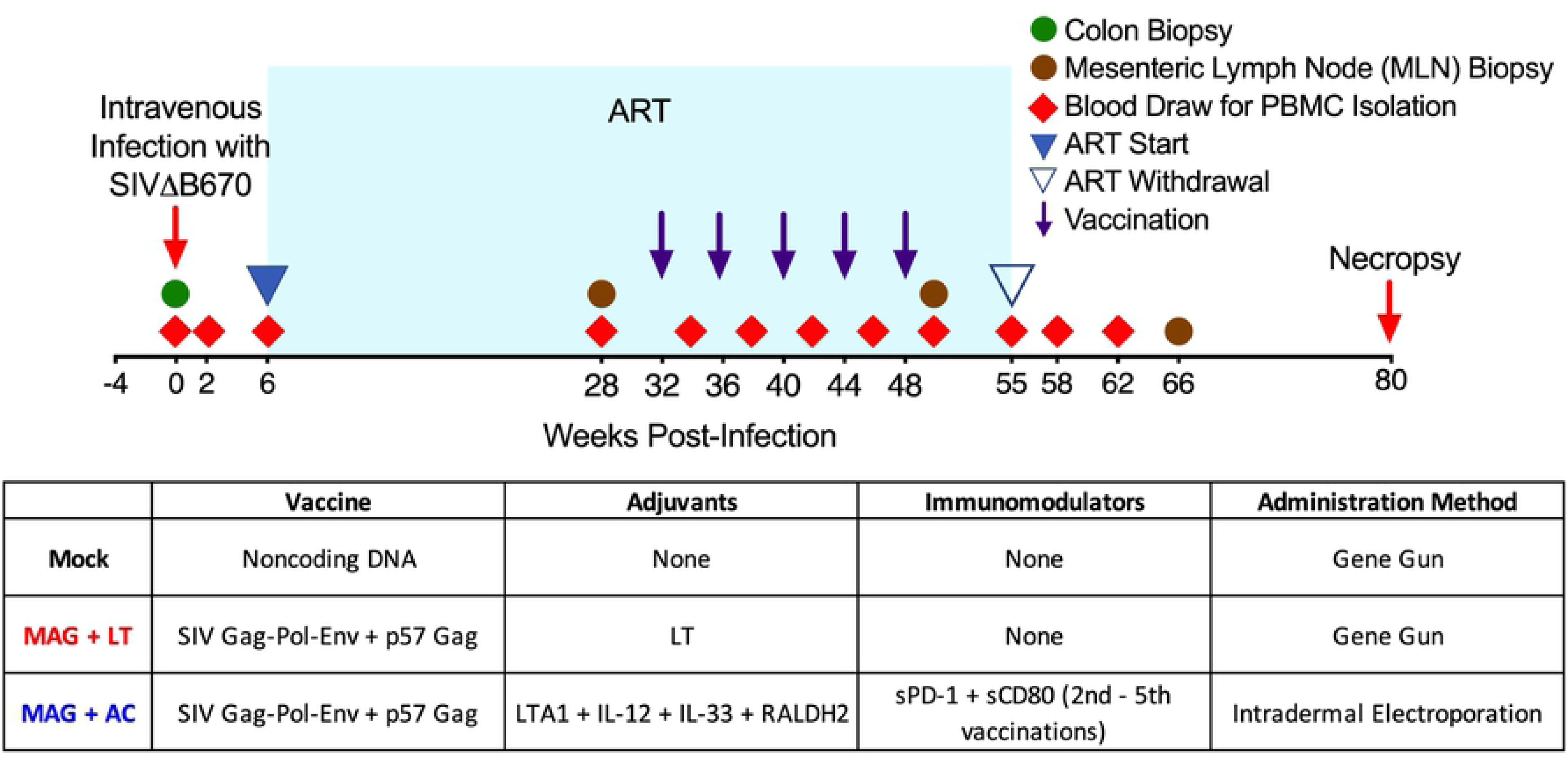
Therapeutic Vaccine Study Design & Response to Antiretroviral Therapy. Indian origin rhesus macaques were infected with SIVΔB670 at week 0 (red arrow) and were treated with ART starting at 6 weeks post-infection (wpi). Purple arrows indicate a series of 5 DNA immunizations spaced 1 month apart, occurring between 32 wpi and 48 wpi. At week 55, ART was interrupted to assess the efficacy of the therapeutic vaccine on viral control. Animals were necropsied at 80 wpi or earlier in the presence of AIDS-defining conditions. Red triangles indicate blood draws for PBMC isolation and brown circles indicate MLN biopsies to measure systemic and gut-associated immune responses. Prior to administering therapeutic immunizations, macaques were stratified so that each group had comparable viral loads and CD4 counts prior to and during ART.

The control group (N = 4) received mock DNA immunizations via particle-mediated epidermal delivery (PMED) consisting of the vaccine plasmid backbone, without SIV antigens or adjuvants. The MAG + LT group (N = 5) received the MAG vaccine, a plasmid that encodes Gag-Pol-Env and expresses virus-like particles as described previously [37] co-delivered with plasmids encoding p57 Gag and the LT adjuvant that we previously showed induces mucosal T cell responses [15, 25], also via PMED. The MAG + AC group (N = 5) received DNA immunizations via intradermal injection followed by electroporation. For the first vaccination, animals received the MAG DNA vaccine co-formulated with plasmids expressing the adjuvants LTA1, IL-12, IL-33, RALDH2. For the subsequent four vaccinations, each macaque in this group also received a plasmid co-expressing sPD-1 and sCD80 in addition to the plasmids encoding the adjuvant combination.

### ART reduces viremia and restores CD4 T cells in the periphery

We and others have shown that therapeutic vaccines are not effective in animals that respond poorly to ART [15, 38]. In addition, our lab and others have shown that, even when HIV or SIV is effectively controlled by ART, increased immune dysfunction, regulatory responses and T cell exhaustion occur that can suppress virus-specific immune responses [39, 40]. To provide potent suppresion of viral replication during ART, we employed a combination of three antiretrovirals: the integrase inhibitor, Raltegravir, and the non-nucleoside reverse transcriptase inhibitors FTC and PMPA that were previously shown to effectively suppress SIV infection in rhesus macaques [41]. However, impaired kidney function due to prolonged treatment with PMPA necessitated a swich over to tenofovir dispoproxil fumarate (TDF) in four animals: A16144 and A16145 in the MAG + LT group, A16149 in the mock vaccine group, and A16237 in the MAG + AC group.

Prior to starting the immunizations at week 32, 12/14 animals responded well to ART, reaching viral loads ≤ 10^3^ viral copies per mL of plasma and resulting in a significant reduction in viral replication (P < 0.0001, S1 Fig). The ART regimen was also effective at restoring blood CD4^+^ T cell counts. During acute infection, animals experienced a rapid CD4^+^ T cell decline that was significantly restored after initiation of ART (P = 0.040, S1 Fig). Overall, there was no significant differences between groups in viral load or peripheral CD4^+^ T cell counts during acute infection and after ART initiation. Two animals exhibited a more modest response to ART, but still showed a 10^4^ decrease in their plasma viremia (S1 Fig) and maintained good health and did not experience disease progression prior to ATI.

### Therapeutic vaccination with MAG + AC increases SIV-specific humoral and cellular responses

To assess the impact of therapeutic vaccination on antibody responses, levels of SIV gp130-specific IgG were measured by ELISA. Peak antibody titers developed after the first vaccine dose (34 wpi) in the MAG + LT animals and after the second dose (38 wpi) in the MAG + AC group, but antibody responses declined by 50 wpi despite the administration of additonal vaccine doses (Fig 2A). Peak antibody responses were significantly higher in the MAG + LT group (P = 0.027, Fig 2B) and trended higher in the MAG + AC group (P = 0.085, Fig 2B) when compared to the mock-vaccinated group, although there was no difference in peak antibody titer between the MAG + LT and MAG + AC vaccinated groups (Fig 2B). Studies in HIV+ people showed strong correlations between HIV-specific antibody-dependent cell-mediated cytotoxicity (ADCC) with protection from, or delayed progression of, HIV infection [42, 43]. To determine if our adjuvanted MAG DNA vaccine elicited antibodies that could mediate ADCC, we analyzed SIV gp130-specific, FcγRIIIA binding antibodies as a marker for antibody dependent cellular cytoxicity (ADCC) [44] by ELISA after the final DNA vaccine dose (50 wpi). The results in Figure 2C showed no significant differences in FcγRIIIA-binding antibody between any of the groups, indicating that in this setting, the therapeutic vaccines likely had no impact on ADCC activity.

**Fig 2.**
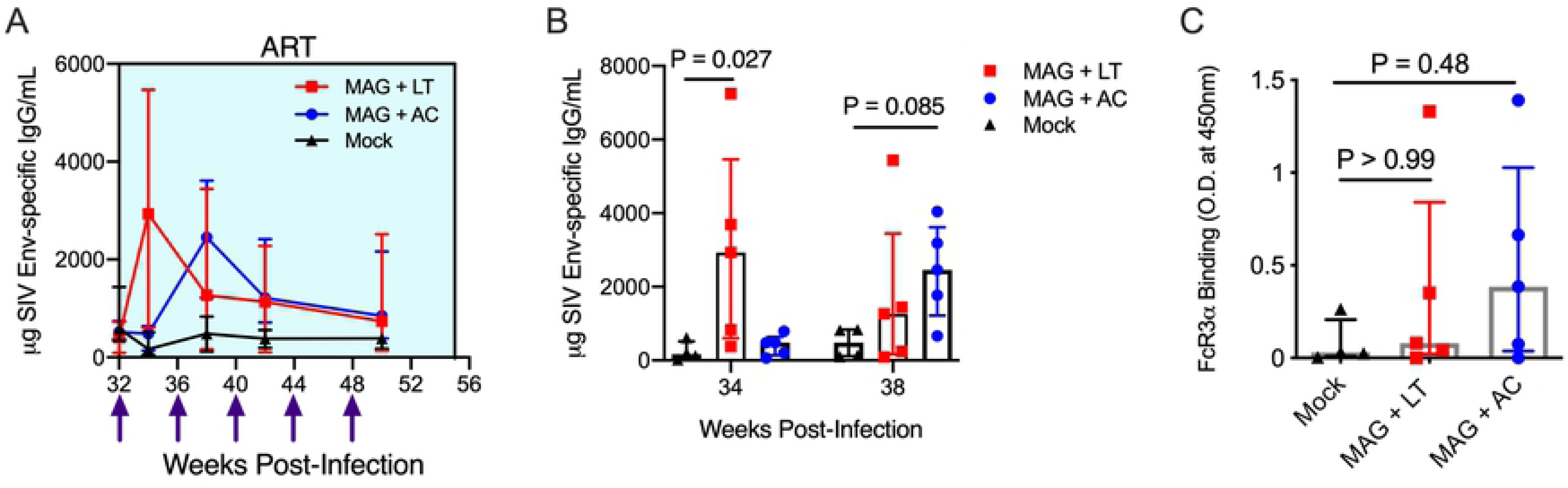
DNA vaccine and adjuvant combinations increase Env-specific antibody responses during ART. **(A)** The magnitude of the SIV Env-specific IgG response in the plasma was measured by ELISA, using SIV gp130 as the capture antigen. Shown are medians and interquartile ranges. **(B)** Peak SIV Env-specific antibody titers are shown for the MAG + LT group (34 wpi) and for the MAG + AC group (38 wpi). Shown are medians and interquartile ranges with individual responses layered over each bar. **(C)** SIV gp130-specific IgG that bind to Fc*γ*R3a, as a surrogate marker for ADCC, was measured using a modified ELISA that employed Fc*γ*R3a conjugated to Streptavidin poly-HRP for the detection antibody. Shown are medians and interquartile ranges with individual responses layered over each bar. **(B, C) Statistics.** A Dunn’s multiple comparisons test was used when making multiple comparisons between vaccine groups and the mock group. Results are considered significant if P ≤ 0.05.

To determine the impact of the therapeutic vaccines on T cell responses, SIV-specific T cells producing IFN-γ in response to stimulation with peptide pools spanning p57^Gag^ (Gag) and gp130 (Env) were measured by ELISpot. The median SIV-specific T cell responses in the MAG+ LT group peaked after the first vaccine dose but steadily declined despite additional vaccine doses (Fig 3A). In contrast, the median T cell responses in the MAG + AC animals increased with each dose up to the third dose (Fig 3A), resulting in significantly higher responses compared to the MAG + LT group at 50 wpi, a timepoint that corresponds to two weeks after the final (5^th^) DNA vaccine dose (P = 0.022, Fig 3B).

**Fig 3.**
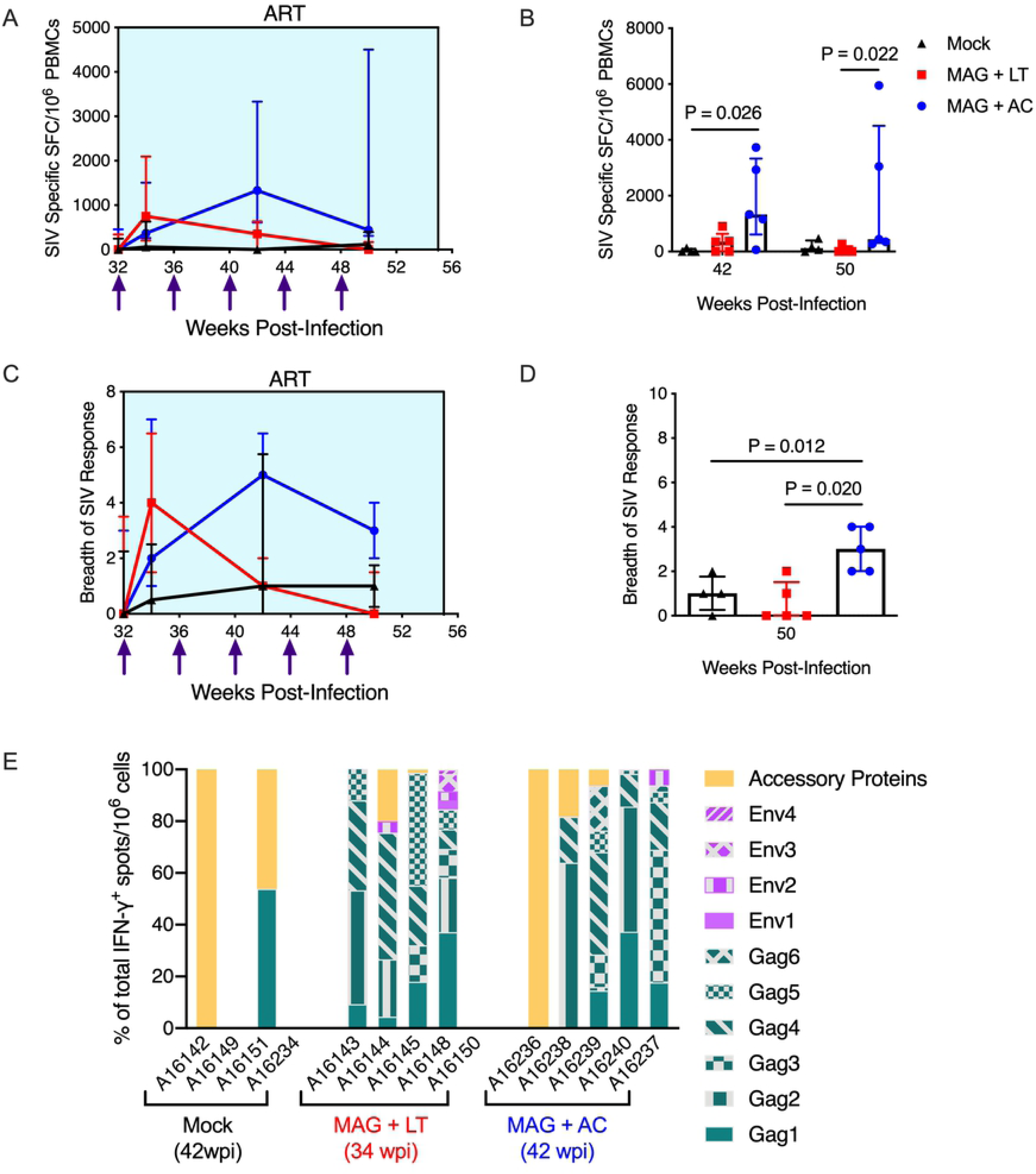
DNA vaccine and adjuvant combinations increase SIV-specific IFN-*γ* T cell responses during ART. **(A-B)** PBMCs were stimulated with Gag, Env, Pol, Vif, Vpr, Rev, Nef, and Tat peptides to quantify the SIV-specific IFN-γ response. Samples were considered positive if peptide-specific responses were at least twice that of the negative control plus at least 0.01% after background (DMSO) subtraction. **(A)** Shown are medians and interquartile ranges of the cumulative (sum of response against all peptides) IFN-γ response. **(B)** The cumulative SIV-specific IFN-γ response is shown after 3 vaccinations (42 wpi) and post-vaccination (50 wpi). Shown are medians and interquartile ranges with individual responses layered over each bar. **(C-D)** The breadth of the SIV-specific IFN-γ response is the number of peptide pools with a positive IFN-γ response. **(C)** Shown are the medians and individual responses layered over each timepoint. **(D)** The cumulative breadth of the the SIV-specific IFN-γ response post-vaccination (50 wpi) is shown. Depicted are medians and interquartile ranges with individual responses layered over each bar. **(E)** The percent of the IFN-γ response specific for Gag, Env and accessory proteins was calculated from the cumulative IFN-γ response at peak breadth (34 wpi for MAG + LT, 42 wpi for MAG + AC and Mock.) **(B, D) Statistics.** A Dunn’s multiple comparisons test was used when making multiple comparisons between vaccine groups and the mock group. Results are considered significant if P ≤ 0.05.

Both vaccines broadened the SIV-specific IFN-γ response after the first dose (34 wpi) when compared to responses measured prior to vaccination or to the control group (Fig 3C). However, likely due to the small group sizes, the increases at this timepoint fell short of statistical significance. Following the final vaccine dose (50 wpi), the breadth of the T cell response had declined in both groups (Fig 3C), although the MAG + AC vaccine group sustained greater breadth than both the MAG + LT group (P = 0.020, Fig 3D) and the mock-vaccinated group (P = 0.012, Fig 3D). Interestingly, the peak IFN-γ T cell response post-vaccination was predominantly directed towards Gag in both vaccine groups, with up to 100% of the IFN-γ response targeting Gag sequences in both the MAG + LT and MAG + AC groups, compared to a maximum of 53% in a single mock-vaccinated animal (Fig 3E).

Env- and Gag-specific CD4^+^ and CD8^+^ T cell responses were further characterized in the PBMC and MLN for effector functions, including secretion of IFN-γ, TNF-α, and IL-2, and co-expression of CD107a/GranzymeB as markers of cytolytic function by intracellular cytokine staining (ICS) and flow cytometry (Fig 4A-B).

**Fig 4.**
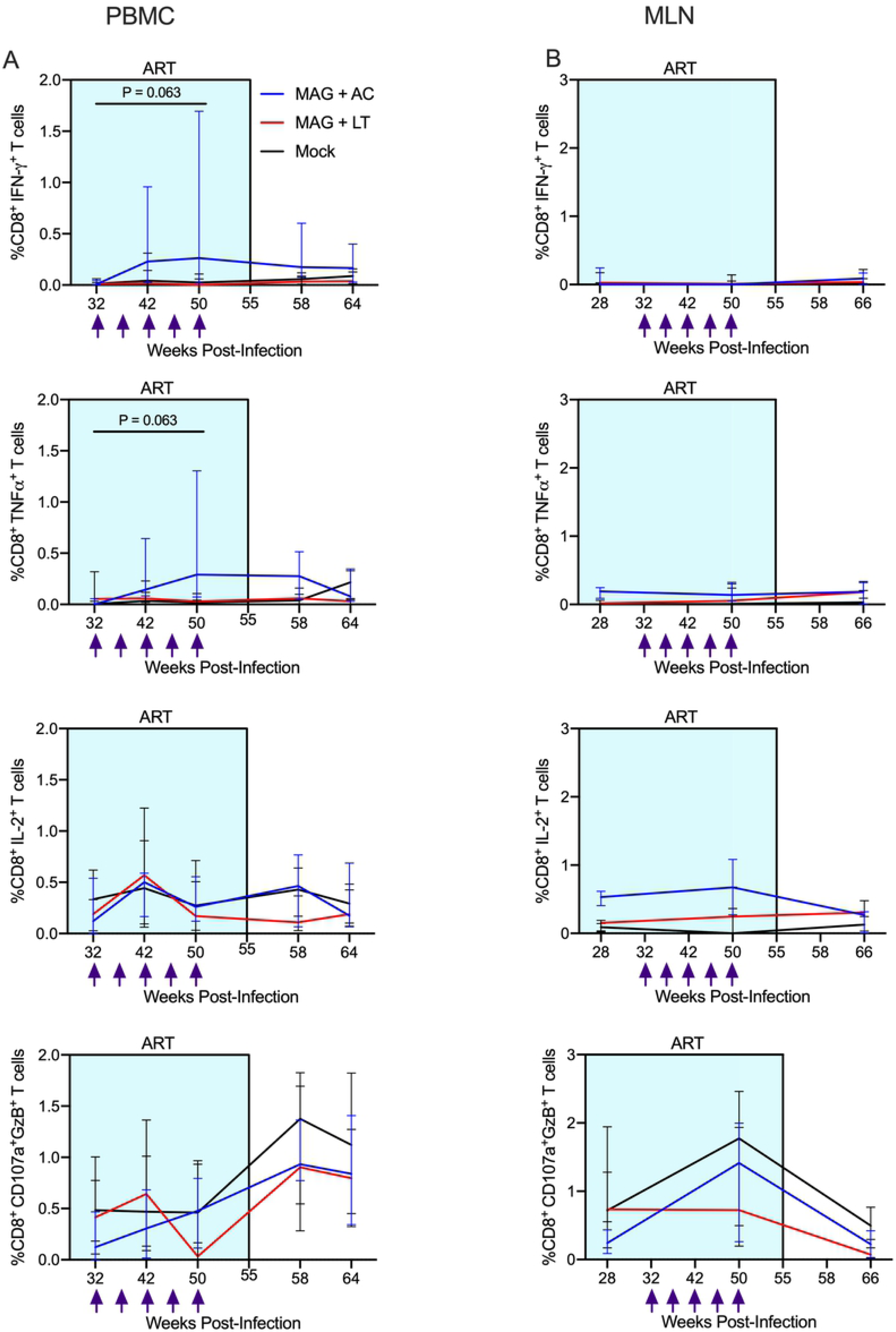
The MAG + AC group shows a trend towards increased IFN-γ and TNFα CD8^+^ T cell responses post-vaccination compared to pre-vaccination. **(A)** PBMCs were thawed and stimulated with Gag peptides, and expression of IL-2, IFN-γ, TNFα and CD107a/GzB were quantified using intracellular cytokine staining. Shown are the medians and interquartile ranges of each group’s SIV-Gag specific T cell response. **(B)** Lymphocytes isolated from MLN were thawed and stimulated with Gag peptides, and expression of IL-2, IFN-γ, TNFα and CD107a/GzB were quantified using intracellular cytokine staining. Shown are the medians and interquartile ranges of each group’s SIV-Gag specific T cell response. **(A, B) Statistics**. Statistical comparisons between baseline and post-vaccination timepoints within a group were calculated using a Wilcoxon matched-pairs signed rank test. Results are considered significant if P ≤ 0.05.

Notably, SIV Gag-specific IFN-γ^+^ and TNF-α^+^ CD8^+^ T cell responses transiently increased after the final vaccination (50 wpi) in only the MAG + AC group, as compared to pre-vaccination baseline levels (32 wpi) (P = 0.063, Fig 4A), although this trend was not sustained during ATI (P > 0.99). Throughout the study, there were no significant differences in Gag-specific CD4^+^ (S2 Fig) or CD8^+^ T cell responses in the MLN between any of the groups (Fig 4B). In addition, frequencies of SIV Env-specific CD8^+^ and CD4^+^ T cell responses in both PBMC (S3 Fig) and MLN (S4 Fig) were similar throughout the study in all three groups.

### Impact of therapeutic vaccination on protection from viral rebound during analytical treatment interruption (ATI)

To determine if therapeutic vaccination improved viral control, ATI was initiated three weeks after the final vaccine dose (55 wpi), and viral loads were monitored for 6 months. Containment of median viral loads at or below 1000 copies/mL of plasma during ATI was chosen as the primary criterion for therapeutic efficacy. This threshold is based on previous studies showing that rhesus macaques infected with SIVΔB670 that maintained viral loads below this level consistently exhibit long term (>1 year) protection from progression to AIDS [37]. Prior to stopping ART, all but one animal in the MAG + AC group (A16239) had viral loads of <1000 RNA copies/mL of plasma (Fig 5A-C). During ATI, 60% (3/5) of animals in the MAG + AC group maintained viremia below 10^3^ RNA copies/mL of plasma for 6 months and sustained CD4 counts above 50% of pre-infection levels, whereas only one animal in the MAG + LT group (1/5, 20%) and one in the control group (1/4, 25%) exhibited similar viral control and protection from disease progression (Fig 5A-C). However, due to the low numbers of animals in each group, these differences were not statistically significant. Vaccinated and control animals that exhibited immediate viral rebound during ATI also showed significant CD4^+^ T cell decline during ATI (S5 Fig). Overall, there was no significant difference in protection from viral rebound or mean viral loads during ATI between the three groups (Fig 5D).

**Fig 5.**
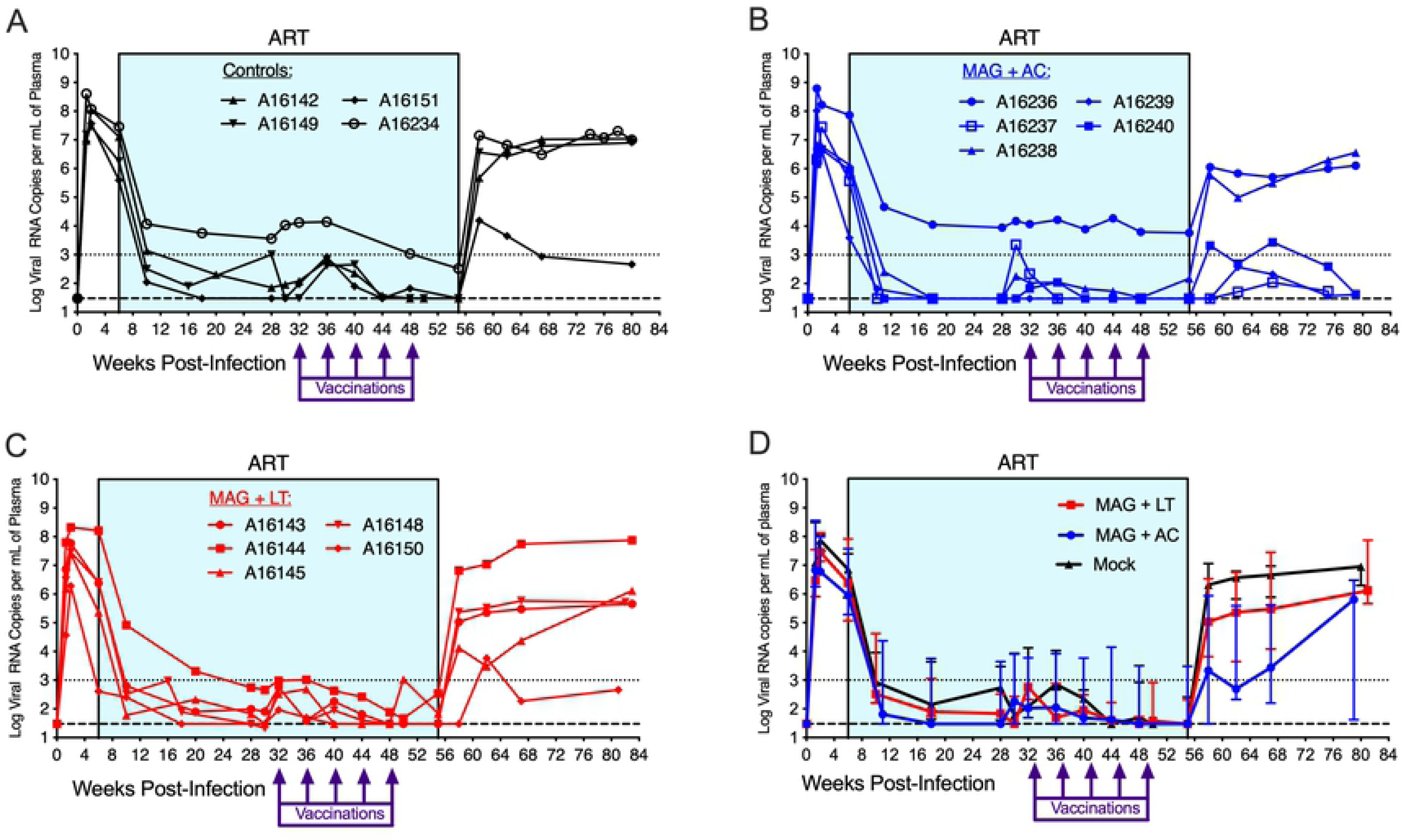
Three out of five animals in the MAG + AC group control virus replication during ATI. **(A-C)** Plasma viral RNA levels were quantified using RT-q-PCR, with a limit of detection of 30 viral RNA copies per 1 mL of plasma, as indicated by the dashed line. **(D)** Shown are the median viral load and interquartile ranges for each treatment group. The dotted line indicates the threshold for control of virus replication, based on previous studies using SIVΔB670.

### Viral control during ATI is associated with increased Gag-specific CD4 and CD8 T cells in MLN and PBMC

The variability in viral rebound and viremia during ATI among all animals in this study enabled further study of immune correlates of viral control. Altogether, within all groups there were 5 viral controllers and 9 noncontrollers, defined as animals that maintained median viremia at or below 1000 copies/mL of plasma or greather than 1000 copies/mL of plasma, respectively, for 5 months after stopping ART (Fig 6A). Viral burden during ATI was significantly different between controllers and noncontrollers (P = 0.0010, Fig 6B).

**Fig 6.**
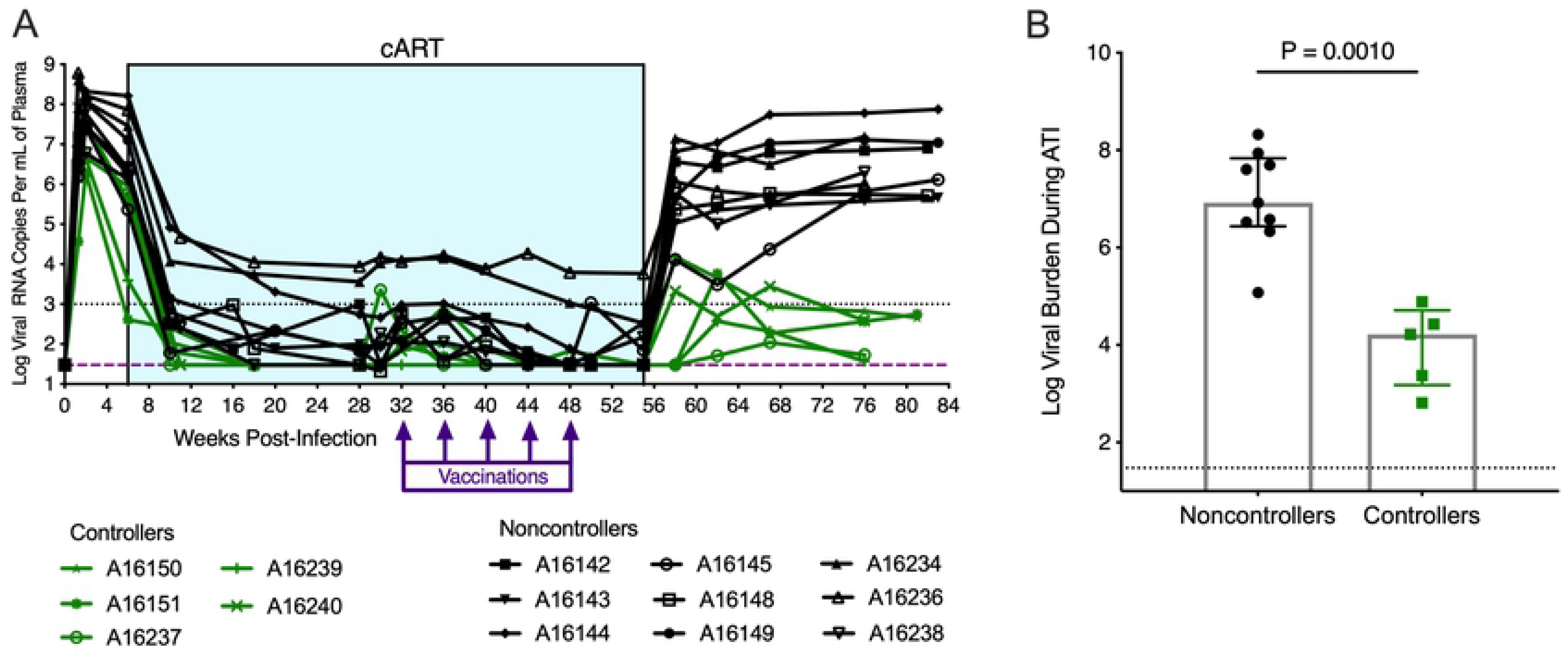
Five controllers maintained significantly lower viral burden during ATI compared to nine noncontrollers. **(A)** Plasma viral RNA levels were quantified using RT-q-PCR, with a limit of detection of 30 viral RNA copies per 1 mL of plasma, as indicated by the dashed line. The dotted line denotes the threshold for control of virus replication, based on previous studies using SIVΔB670. Controllers were defined as animals that maintained a median viremia at or below 1000 viral RNA copies per 1 mL of plasma for 5 months post-ART. Noncontrollers were defined as animals with a median viremia that exceeded 1000 viral RNA copies per 1mL of plasma for 5 months post-ART. **(B)** Viral burden during ATI was calculated as the area under the curve of each animal’s viral load from 55 wpi to 76 wpi, shown are the median viral burden and interquartile ranges. Statistics were calculated using a Mann-Whitney t test. Results are considered significant if P ≤ 0.05.

To determine immune correlates of viral control, we compared frequencies of Gag-specific CD4^+^ and CD8^+^ T cell responses expressing IFN-γ, TNFα, IL-2, and/or co-expressing the cytolytic markers CD107a/GranzymeB as detected by flow cytometry in the controllers and noncontrollers. Immune responses were compared at key timepoints: After the final vaccination and prior to ATI (50 wpi for both PBMC and MLN) and after viral setpoint was established (62 wpi for PBMC and 66 wpi for MLN). In PBMC, controllers demonstrated a trend towards higher frequencies of TNFα^+^ CD8^+^ T cells prior to ATI (50 wpi) (Fig 7A, P = 0.056). Controllers also exhibited significantly increased frequencies of Gag-specific IFN-γ^+^ CD8^+^ T cells in PBMC during ATI (62 wpi) (Fig 7B, P = 0.0080). However, we did not observe any differences between controllers and noncontrollers in frequencies of Gag-specific IL-2^+^ CD8^+^ or CD107a^+^/GzB^+^ CD8^+^ T cells in PBMC either prior to or during ATI (S5 Fig, S6 Fig). In MLN, controllers showed significantly increased frequencies of Gag-specific IL-2^+^ CD8^+^ T cells when compared to noncontrollers (Fig 7C, P = 0.037), although these differences were not sustained during ATI (66 wpi). Additionally, there were no differences between controllers and noncontrollers in Gag-specific CD8+ T cells in MLN expressing IFN-γ, TNFα, or co-expressing CD107a^+^/GzB^+^ (S5 Fig, S6 Fig).

**Fig 7.**
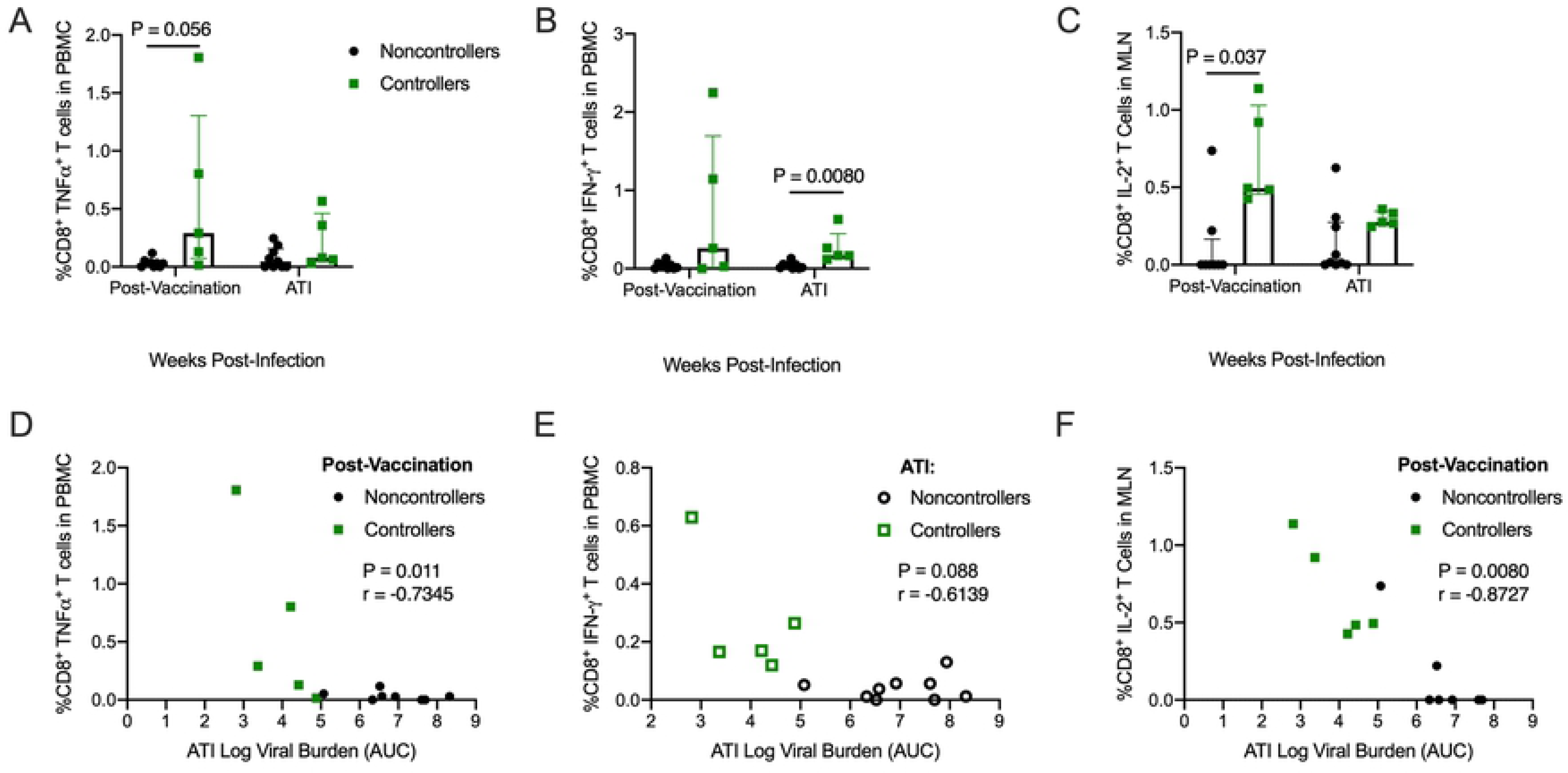
Controllers have higher SIV Gag-specific CD8^+^ T cell responses in PBMC and MLN post-vaccination and during ATI. **(A-C)** PBMCs and MLNs were thawed and stimulated with Gag peptides, and expression of cytokines was quantified using intracellular cytokine staining. Shown are the medians and interquartile ranges of the SIV Gag-specific CD8^+^ T cell responses of controllers and noncontrollers, with individual responses layered over each bar at a post-vaccination timepoint (50 wpi) and during ATI (62 wpi for PBMC and 66 wpi for MLN). Statistical differences between controllers and noncontrollers at each timepoint were calculated using a Mann Whitney t test. Benjamini-Hochberg adjusted P values are shown, results are considered significant if P ≤ 0.05. **(D)** The SIV Gag-specific TNFα CD8^+^ T cell responses in PBMC at 50 wpi negatively correlated with the viral burden measured as area under the curve (AUC) during ATI. **(E)** The SIV Gag-specific IFN-γ CD8^+^ T cell responses in PBMC at 50 wpi negatively correlated with the viral burden measured as area under the curve (AUC) during ATI. **(F)** The SIV Gag-specific IL-2 CD8^+^ T cell responses in MLN at 62 wpi negatively correlated with the viral burden measured as area under the curve (AUC) during ATI. The P and r values shown were calculated using a Spearman rank correlation test. Benjamini-Hochberg adjusted P values are shown, results are considered significant if P ≤ 0.05.

Importantly, higher frequencies of Gag-specific IFN-γ^+^ and TNFα^+^ CD8^+^ T cell responses in PBMC and Gag-specific IL-2^+^ CD8^+^ T cell responses in the MLN significantly correlated with lower viral burden during ATI (Fig 7D-F), suggesting that SIV Gag-specific T cell responses in both the periphery and GALT may contribute to the improved control of virus replication in controllers.

In contrast, we observed no correlations between Gag-specific CD4^+^ T cell responses in PBMC or MLN and viral control (S7 Fig). We also observed that noncontrollers exhibited a consistent trend towards higher titers of Env-specific IgG both during ART treatment and during ATI (S8 Fig), implying that these antibody responses were likely driven by virus replication and that viral control during ATI was primarily mediated by CD8^+^ T cell responses.

### Viral control during ATI is associated with increased polyfunctionality in MLN and PBMC

To further elucidate the role of SIV-specific CD8^+^ T cell responses in viral control, we next compared the magnitude of the polyfunctional CD8^+^ T cell response, as defined by the frequency of SIV-specific cells specific for either Gag or Env and expressing any three or more of the cytokines IFN-γ, TNFα, IL-2, and/or co-expression of the cytolytic markers CD107a/GranzymeB. Post-vaccination (50 wpi), controllers demonstrated a trend towards higher frequencies of polyfunctional CD8^+^ T cells expressing three effector functions in the PBMC (P = 0.13, Fig 8A) and MLN (P = 0.18, Fig 8B) compared to noncontrollers. During ATI (62 wpi in PBMC and 66 wpi in MLN), controllers exhibited significantly increased frequencies of polyfunctional CD8^+^ T cells in PBMC (P = 0.036, Fig 8A) and a continued trend towards increased frequencies of polyfunctional CD8^+^ T cells in MLN (P = 0.084, Fig 8B) when compared to noncontrollers. Furthermore, higher frequencies of polyfunctional CD8^+^ T cell responses in both the PBMC and MLN correlated with lower viral burden during ATI (Fig 8C-F). These results indicate that polyfunctional CD8^+^ T cell responses localized in both the periphery and gut likely played an integral role in controlling viral recrudescence and protection from disease progression during ATI.

**Fig 8.**
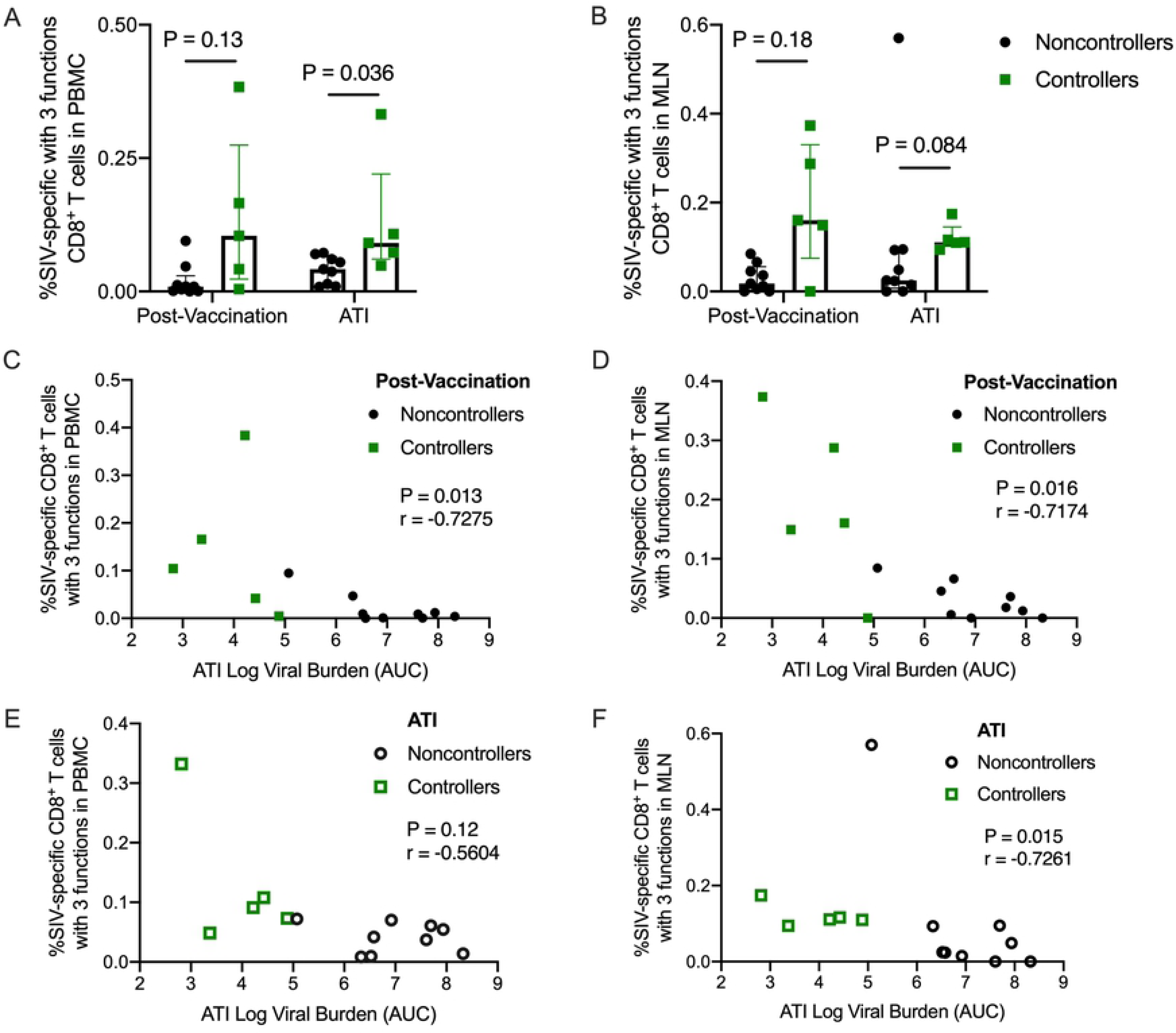
Controllers demonstrate increased frequencies of polyfunctional CD8^+^ T cells in MLN and PBMC. **(A-B)** For each animal, the frequencies of SIV-specific CD8^+^ T cells expressing any 3 combinations of effector functions were summed up. Shown are the medians and interquartile ranges of polyfunctional SIV-specific CD8^+^ T cells, with individual values layered over each bar. Statistical differences between controllers and noncontrollers at each timepoint were determined using a Mann Whitney t test. Benjamini-Hochberg adjusted P values are shown. Results are considered significant if P ≤ 0.05. **(C-D)** The polyfunctional SIV-specific CD8^+^ T cell responses post-vaccination and during ATI negatively correlated with ATI viral burden. The P and r values shown were calculated using a Spearman rank correlation test. Benjamini-Hochberg adjusted P values are shown, and results are considered significant if P ≤ 0.05.

We previously showed that viral control following therapeutic vaccination occurred only in animals that developed very low to undetectable viremia during ART treatment [15, 38]. To determine if acute viral replication or the response to ART influenced viral control during ATI, we compared viral loads measured during acute infection (0-6 wpi) or during ART (6-55 wpi) in controllers and noncontrollers. Controllers demonstrated a trend toward lower viral burden during acute infection (P = 0.11, Fig 9A) and had significantly lower viral burden while on ART (P = 0.014, Fig 9A) when compared to noncontrollers. Furthermore, the level of viral control during ATI significantly correlated with viral burden during acute infection (P = 0.0078, Fig 9B) and during ART treatment (P = 0.00040, Fig 9C). Together, these data suggested that baseline factors influence viral replication during acute infection, that in turn determines an SIV-infected animal’s ability to respond to ART. ART responsiveness then potentially affects the ability of the SIV-infected animals to develop polyfunctional CD8^+^ T cell responses and control viral replication during ATI.

**Fig 9.**
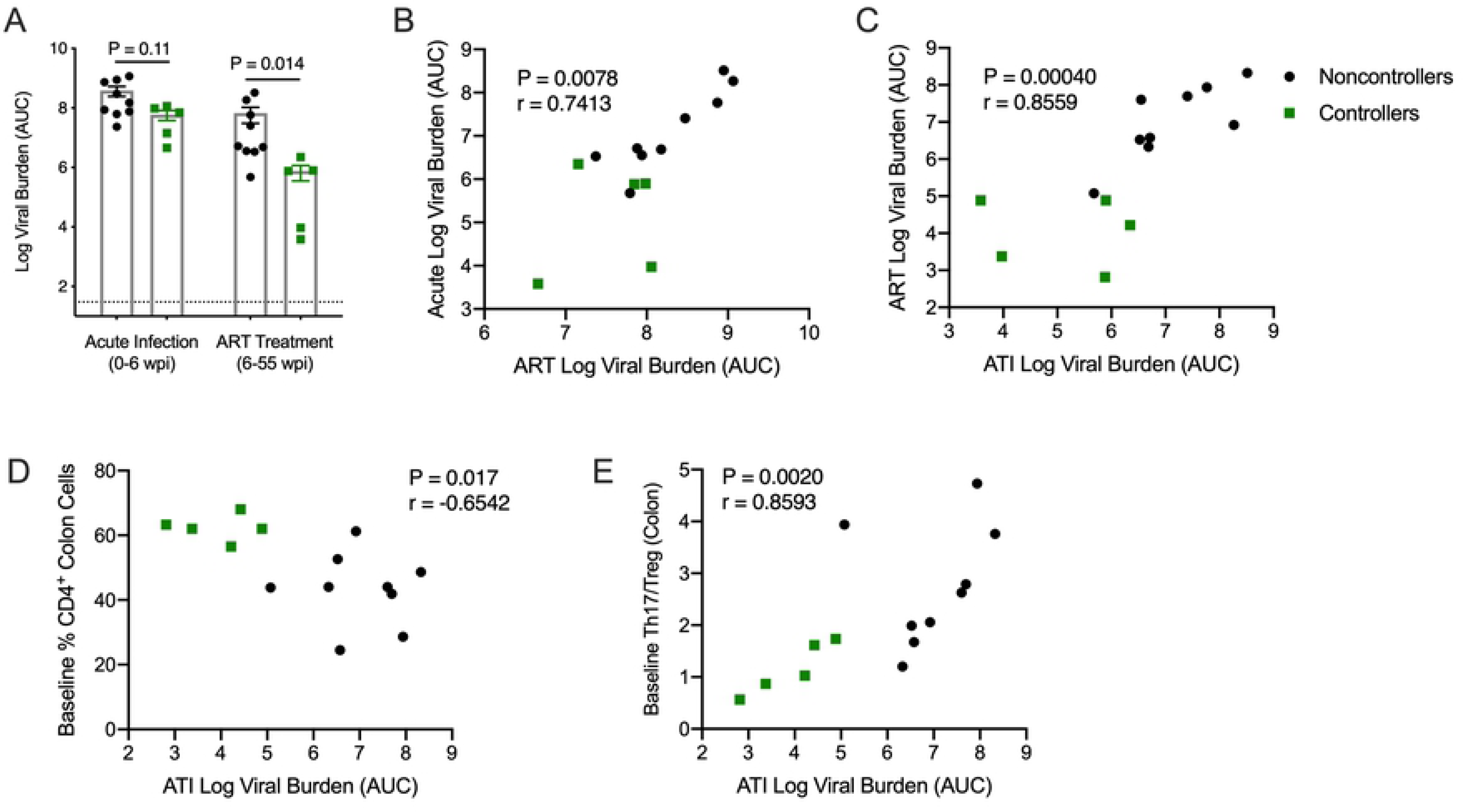
Lower viral replication during acute infection correlates with improved ART responsiveness that in turn predicts therapeutic outcome. **(A-C)** Viral loads were measured via RT-q-PCR and viral burden was calculated as the area under the curve of animals’ viral loads. **(A)** Statistics were calculated using a Mann-Whitney t test. Benjamini-Hochberg adjusted P values are shown. Results are considered significant if P ≤ 0.05. **(B-C)** Acute log viral burden (0 – 6 wpi) correlated with log viral burden during ART treatment (6 – 55 wpi). ART log viral burden in turn correlated with log viral burden during ATI (55 wpi – 76 wpi). **(D)** The frequency of CD4^+^ T cells in the colon at baseline (0 wpi) correlated with viral burden during ATI. **(E)** The ratio of Th17 and Treg cells at baseline correlated with viral burden during ATI. **(B-E)** A Spearman rank correlation test was used to determine P and r values. Shown are Benjamini-Hochberg adjusted P values, results are considered significant if P ≤ 0.05.

To investigate this theory, we assessed a number of baseline immune factors, including CD4^+^ T cells, T helper 17 (Th17) and T regulatory (Treg) cells in the colon, that we hypothesized could influence the extent of acute viral replication. In particular, preferential depletion of Th17 cells in the gut during acute infection corresponds to a loss of gut integrity, leading to microbial translocation, immune activation, and disease progression [45, 46]. In contrast, the role of Treg cells in HIV pathogenesis is not well characterized, but they may contribute to viral replication by suppressing HIV-specific CD8^+^ T cell activity [47]. Alternatively, Tregs may decrease chronic immune activation, thus slowing disease progession [48]. Ultimately, their role in HIV disease progression may depend on the stage of infection and the relative proportion of other T cell subsets. In the colon, we observed a correlation between greater CD4^+^ T cell frequencies (P = 0.017) or lower Th17/Treg ratios (P = 0.0020) with lower viral burden during ATI (Fig 9D-E). Upon further examination, neither the baseline frequencies of Th17 (P = 0.22) nor Treg (P = 0.16) cells alone correlated with viral burden during ATI (S9 Fig). Collectively, this data suggests that during the early stages of infection mucosal T cells are necessary to maintain gut homeostasis and prevent immune dysfunction.

## Discussion

The primary goal of this study was to determine if a multi-antigen DNA vaccine (MAG) that we previously showed imparted viral control when tested as a prophylactic DNA vaccine against SIV infection [37] could be employed as a therapeutic vaccine, and if a novel genetic adjuvant combination (AC) could improve immunogencity and therapeutic efficacy of this vaccine. In support of this, animals that received the multi-antigen vaccine and adjuvant combination (MAG + AC) exhibited significant increases in the magnitude and breadth of IFN-γ T cell responses as measured by ELISpot when compared to a vaccine group that received MAG with only a single adjuvant (LT). Additionally, the MAG + AC group exhibited a trend towards elevated Gag-specific TNFα and IFN-γ CD8^+^ T cell responses in the blood post-vaccination, an outcome that is consistent with our prophylactic vaccine study employing the same vaccine [37] and other vaccine trials in NHP and mice using the IL-12 and IL-33 adjuvants [26, 29, 49]. However, the addition of the RALDH2 adjuvant to the combination did not increase SIV-specific T cell responses in the GALT, as it did in mice [33], possibly due to SIV infection causing significant mucosal immune dysfunction that may have interfered with the effects of this adjuvant. Three out of five MAG + AC animals were protected from viral rebound (60%), although due to the small group sizes, this outcome was not statistically different from the controls. Furthermore, since we were unable to include a control group receiving the full adjuvant combination in the absence of a vaccine, we cannot rule out the possibility that the combined effects of the adjuvant combination may have stimulated nonspecific immune responses that contributed to improved viral control during ATI in these three animals, similar to what was observed by Sui *et al.* in 2010 [50]. We additionally were unable to include a control group to account for the potential differences in MAG immunogenicity resulting from vaccine administration using Gene Gun versus intradermal electroporation. Both vaccine modalities have been shown to enhance DNA vaccine immunogenicity [51, 52], but we cannot rule out that the differences between the methods of vaccine administration may have influenced immunogenicity between the MAG + LT group and the MAG + AC group.

Overall, five animals from all groups controlled viral rebound and were protected from progression to AIDS, in contrast to nine animals that exhibited immediate viral rebound during ATI. This variability in viral control during ATI enabled analysis of immune correlates of protection from viral rebound. We found that polyfunctional, SIV-specific CD8^+^ T cells in the MLN measured prior to stopping ART inversely correlated with viral loads during ATI, a finding that supports our previous study where we found that broadly specific IFN-γ T cell responses localized in the gut, but not the blood, significantly correlated with protection from viral rebound [15]. Notably, although we and other groups in addition to ours have demonstrated that robust, virus-specific CD8^+^ T cell responses in the blood are associated with control of viral replication during ATI, and it is well known that polyfunctional, virus-specific CD8 T cells in the blood are associated with elite control of HIV and SIV, this study is the first to show an association between polyfunctional CD8 T cells in the gut-associated lymphoid tissue (GALT) and control of viral replication during ATI. Together, these studies provide strong evidence that an effective therapeutic vaccine may need to induce not only broadly specific but also polyfunctional mucosal CD8^+^ T cells to effectively control reactivating virus during ATI in this viral reservoir. These data also further corroborate our previous study, where we showed that SIV-specific TNFα^+^ CD107a^+^ CD8^+^ T cells in the blood were associated with protection from viral rebound [15].

While our data clearly show an impact of the vaccines on immune responses and a protective role for mucosal and systemic polyfunctional CD8^+^ T cell responses, the precise mechanisms underlying viral control during ATI in the controllers in this study and the factors that influenced failure to control virus during ATI in the noncontrollers are not clear. An important variable in this study is the relative response to the antiretroviral drug therapy. We previously showed that animals that respond poorly to ART also responded poorly to therapeutic vaccination [15, 38], and our results here showing that the controllers had lower viral loads than the noncontrollers during ART are consistent with these findings. It is notable that the controllers also exhibited a trend toward lower acute viral loads when compared to the noncontrollers, although this was not statistically significant. This suggests that virus-host interactions that occurred prior to ART initiation may have impacted ART efficiency which, in turn, altered the efficacy of therapeutic vaccination. Indeed, what occurs during acute infection, particularly in the mucosa, may determine how well an animal’s viral loads are controlled by ART, thereby influencing the response to therapeutic vaccination and its ability to enhance immune control of viremia during ATI. In support of this, we showed that lower frequencies of colonic CD4^+^ T cells and higher ratios of Th17 to Treg cells pre-infection correlated with higher viral burden during ATI. This could indicate that although Th17 cells are important for maintenance of gut barrier function, they could act as early viral targets of infection. Additionally and alternatively, lower proportions of immunosuppressive Treg cells in the gut mucosa may allow for greater T-cell proliferation and promote early viral replication in the GALT. Meanwhile, having an larger overall population of mucosal CD4^+^ T cells at baseline could indicate that an animal is better able to cope with CD4^+^ T cell depletion and may experience less immune dysfunction as a result of viral replication. We also previously showed that SIV-infected rhesus macaques that were unable to maintain their mucosal Th17/Treg ratios during acute infection and prior to ART initiation, exhibited a significantly lower response to ART [38]. This suggests that disruption of mucosal Th17 and Treg homeostasis during acute infection could also impede an animal’s ability to respond to subsequent therapeutic interventions.

In summary, the significant correlation between higher polyfunctional CD8^+^ T cells in the MLN and lower viral burden during ATI shown here provides new insight into a role for potent, polyfunctional CD8^+^ T cell responses in the GALT in controlling viral rebound, and provides further evidence supporting development of therapeutic HIV vaccines that can induce mucosal immunity. Although we did not observe a significant impact of the MAG + AC vaccine on mucosal immunity, the vaccine effectively increased the magnitude and breadth of the IFN-γ T cell response in the blood, suggesting that the adjuvant combination could be a useful adjuvant for other T cell-based vaccines where peripheral T cell responses are sufficient for protection. Studies are underway to further refine this adjuvant combination to increase its ability to induce mucosal immune responses for future therapeutic vaccine studies. Toward the goal of developing an effective therapeutic vaccine for HIV, further studies are needed to define additional host factors and immune mechanisms induced during acute infection that influence the relative response to ART and the immunogenicity and efficacy of therapeutic vaccines.

## Methods

### Ethics Statement

Male, adult rhesus macaques (*Macaca mulatta*) of Indian origin were used for this study. These animals were housed at the Washington National Primate Research Center (WaNPRC), which is accredited by the American Association for the Accreditation of Laboratory Animal Care International (AAALAC). At the WaNPRC, animals received the highest standard of care from a team of highly trained, experienced animal technicians, veterinarians, and animal behavior specialists, who provided daily environmental enrichment and monitored for any signs of distress or abnormal behavior. All biopsies, surgeries, and blood draws were performed under ketamine anesthesia and any continuous discomfort or pain was alleviated at the discretion of the veterinary staff. Following SIV infection, animals were monitored for signs of disease progression, including CD4^+^ T cell count, weight, anemia, and opportunistic infections, at least monthly. The University of Washington’s Institutional Animal Care and Use Committee (IACUC) approved all experiments in these macaques.

### MHC-I and TRIM5 typing

All macaques were major histocompatibility class I (MHC-1) typed for 32 alleles, including A*01, A*02, B*08, B*17. DNA was extracted using the Roche© MagnaPure™ system and analyzed via PCR by Dr. David Watkins and the MHC Genotyping Service at the University of Miami, as previously described [53, 54]. All animals were also tested for TRIM5 haplotypes, including TFP, Q, and CypA, by PCR of genomic DNA by Dr. David O’Connor at the Wisconsin National Primate Research Center (WNPRC). Animals with permissive TRIM5 genotypes were excluded from the study, as were animals possessing restrictive TRIM5 genotypes and MHC haplotypes associated with increased viral control.

### Viral Challenge and AIDS Monitoring

Rhesus macaques were challenged intravenously with 100 TCID_50_ of cryopreserved SIVΔB670, diluted in 1 milliliter of RPMI. Simian AIDS was defined according to WaNPRC guidelines, namely: weight loss exceeding 15 percent, anemia, CD4^+^ T cell decline to less than 200 cells per microliter, and presence of opportunistic infections. These criteria were evaluated at least monthly, but if two or more of the criteria were met, these measurements were evaluated more frequently. If veterinary staff determined that an animal had reached AIDS-defining criteria, humane euthanasia was performed as an early endpoint.

### Quantification of Plasma Viral Load and Complete Blood Counts (CBCs) and Serum Chemistries

The Virology Core at the WaNPRC, led by Dr. Shiu-Lok Hu and Dr. Patricia Firpo, quantified viral RNA in the plasma of SIVΔB670-infected animals using a real time quantitative PCR (RT-q-PCR) assay. The Virology Core also determined complete blood counts, using a Beckman Coulter® AC*T^TM^ 5diff hematology analyzer as described previously [55].

### Antiretroviral Therapy

All SIV-infected animals were treated with a combination of 3 antiretroviral therapies: 9-(2-Phosphoryl-methoxypropyly) adenine (PMPA or tenofovir; Gilead Sciences, Foster City, CA) was resuspended in phosphate-buffered saline (PBS) at120 mg/mL. To completely dissolve the PMPA, 1 molar NaOH was added until the pH reached 7.4-7.8. The solution was then filter purified, injected into sterile glass vials, and stored at -20°C. PMPA was administered subcutaneously in a once-daily dose of 20 milligrams per kilogram (mg/kg) of animal weight. 2’,3’-dideoxy-5-fluroro-3’-thiacytidine (FTC or emtricitabine, Gilead Sciences, Foster City, CA) was resuspended in PBS at 120 mg/mL. The mixture was heated at 37°C with constant stirring until completely dissolved, and stored at 4°C. FTC was administered once per-day subcutaneously at 30 mg/kg during the first month of ART (weeks 6-10 post-infection) and at 20 mg/kg once per-day for the remainder of ART. Raltegravir (Isentress, Merck & Co., Kenilworth, NJ) was given orally at 250 mg/animal twice daily for the first month of ART, and at 150 mg/animal twice daily for the remainder of ART.

Trained animal technicians administered all ART drugs, and veterinary staff closely monitored animals for adverse side effects, which were treated immediately at their discretion.

A few animals experienced elevated creatinine levels due to prolonged treatment with PMPA, so these animals were promptly switched to tenofovir disoproxil fumarate (TDF), a prodrug of tenofovir that is metabolized to PMPA.

TDF was resuspended in a solution of 15% Kleptose in water, at a concentration of 10.2 mg/mL, and pH adjusted to 4.1-4.3. The solution was then filter purified and stored at 4°C or frozen at -20°C for long-term storage. TDF was administered once per-day subcutaneously at 5.2 mg/kg for the duration of ART.

### DNA Vaccinations

#### I. Particle-Mediated Epidermal Delivery (PMED, or Gene Gun)

Vaccine and adjuvant plasmids were formulated onto gold particles as previously described, and administered using the PowderJect® XR1 gene delivery device (PowderJect Vaccines, Inc., Middleton, WI) [15]. Fur was shaved off of vaccination sites, which were then swabbed with alcohol prior to vaccine administration. Macaques were vaccinated over 16 epidermal sites along the lower abdomen and over the inguinal lymph nodes. Each animal received 32 μg of the MAG or Gag DNA vaccine co-formulated with 3.2 μg of plasmid expressing the LT adjuvant (2 μg MAG or Gag DNA + 0.2 μg LT per site).

#### **II.** Intradermal Electroporation (ID EP)

MAG, Gag, and adjuvant plasmids (rhIL-12, LTA1, expressed on one plasmid each, and hRALDH2/rhIL-33 and rhPD-1/rhCD80 co-expressed on one plasmid each) were prepared in a citrate buffer and administered via intradermal injection into the dermis above the quadriceps muscle on each leg. For the first vaccination, each macaque received 900 μg of the MAG or Gag DNA vaccine co-formulated with 900 μg of DNA expressing hRALDH2/rhIL-33, 900 μg of DNA expressing rhIL-12, and 162 μg of DNA expressing LTA1, evenly distributed over 3 injection sites per leg (300 μg MAG or Gag + 300 μg hRALDH2/rhIL-33 + 300 μg rhIL-12 + 54 μg LTA1 per site). For each subsequent vaccination, each macaque received 900 μg of the MAG or Gag DNA vaccine co-formulated with 900 μg of DNA expressing hRALDH2/rhIL-33, 900 μg of DNA expressing rhIL-12, 975 μg of DNA expressing rhPD-1/rhCD80 and 162 μg of DNA expressing LTA1, evenly distributed over 4 injection sites per leg (225 μg MAG or Gag + 225 μg hRALDH2/rhIL-33 + 225 μg rhIL-12 + 244 μg rhPD-1/CD80 + 40.5 μg LTA1 per site). Prior to each vaccination, fur covering the vaccination site was shaved and the skin was swabbed with alcohol. Following injection of vaccine and adjuvant DNA, electrical pulses were delivered using the Agile Pulse device (BTX, Holliston, MA) according to the device manufacturer’s instructions.

### Luminex®

To quantify the levels of 23 cytokines in the plasma of SIV-infected animals, the Milliplex® Map Non-Human Primate Cytokine Magnetic Bead Panel Kit (EMD Millipore Corporation, Billerica, MA) was used, according to the manufacturer’s instructions.

### Enzyme-Linked Immunospot Assay (ELISpot)

ELISpot was performed to quantify the frequency of SIV-specific IFN-γ spot-forming cells (SFC) in accordance with previously described methods. In brief, PBMCs were isolated from whole blood via density gradient separation and stimulated with pools of 15mer peptides overlapping by 11 amino acids and corresponding to the following SIV proteins: Gag, Env, Pol, Vif, Vpr, Rev, Nef, and Tat. As a negative control, samples were stimulated with dimethyl sulfoxide (DMSO). For a positive control, samples were stimulated with concanavalin A (Con A). Samples were considered positive if peptide-specific responses were at least twice that of the negative control plus at least 0.01% after background (DMSO) subtraction.

### Enzyme-Linked Immunosorbent Assay (ELISA) for analysis of antibody responses and microbial translocation

SIV Env-specific IgG binding antibody was measured by ELISA, as previously described. In brief, 1μg/mL SIVmac239 gp130 (NIH AIDS Reagent Program) was used as the capture antigen, and a rabbit anti-IgG (heavy and light chains conjugated to horse radish peroxidase) was used to detect antibody bound to the capture antigen.

IgG binding to FcγR3a was measured using a modified ELISA protocol. SIVmac239 gp130 (NIH AIDS Reagent Program) was again used as the capture antigen, and serial dilutions of macaque plasma in Blotto buffer (20x TRIS-buffered saline and 2% Tween20 diluted to 1x with ddH_2_O and 5% non-fat milk) were plated in duplicate and incubated for one hour at room temperature. While the experimental plate was incubating, 24μL of 250μg/mL biotinylated FcγR3a was mixed with 5μL of 0.5mg/mL of Streptavidin poly-HRP in 3.3mL Blotto buffer to bind the HRP probe to the recombinant FcγR3a. This mixture was used in place of a detection antibody to determine the concentration of Env-specific IgG that binds to FcγR3a. The FcγR3a mixture was incubated for one hour at room temperature on a rotator, and then diluted 1:3 with Blotto buffer. Once the FcγR3a mixture was prepared and the experimental plate had finished its one hour incubation, the plate was washed three times with Blotto buffer, and 100μL of the FcγR3a mixture was added. The plate was then incubated once more for one hour, then washed three times with Blotto buffer, developed using the SureBlue™ TMB Microwell Peroxidase Substrate Kit (KPL Inc.) and neutralized after 10 minutes with 1N H_2_SO_4_. The optical density of the samples was measured using an EMax® ELISA Microplate Reader with SoftMax® Pro software (Molecular Devices©, Sunnyvale, California). Samples were background subtracted from negative control wells to which no macaque plasma was added, and a positive response was defined as greater than two standard deviations above the mean OD of the negative control wells.

### Intracellular Cytokine Staining (ICS) and immunophenotyping of T cell exhaustion

Cryopreserved PBMCs and MLN lymphocytes were thawed and rested at 37°C and 5% CO_2_ for 6 hours before stimulation with DMSO, PMA (Sigma-Aldrich®)/Ionomycin (Life Technologies®), or SIV Gag or Env peptides (1 μg/mL) for 1 hour with CD107a PECy5 (eBioH4A3, BioLegend) in R10 media before adding 1 mg/mL of Brefeldin A (Sigma-Aldrich®). Cells were stimulated overnight (approximately 14 hours) at 37°C and 5% CO_2_. After stimulation, cells were washed with PBS and stained using LIVE/DEAD® Aqua (ThermoFisher®) amine reactive dye to distinguish live cells, then surface stained with CD3 Brilliant Violet (BV) 711 (Sp34-2, BD Biosciences), CD4 PerCPCy5.5 (L200, BD Biosciences), CD8 APC-Cy7 (RPA-T8, BD Biosciences), CD28 PE-CF594 (CD28.2, BD Biosciences), CD95 BV421 (Dx2, BD), PD-1 BV605 (EHI12.2H7, BioLegend), and TIGIT PerCP-eFluor710 (MBSA43, ThermoFisher®), in Brilliant Stain buffer (BD Biosciences). Cells were then permeabilized with Cytofix/Cytoperm (BD Biosciences) and stained for intracellular cytokines with an antibody cocktail of IFNγ FITC (B27, BD Biosciences), TNFα PE-Cy8 (Mab11, BD Biosciences), IL-2 PE (MQ1-17H12, BD Biosciences), and GranzymeB APC (GB12, ThermoFisher®), in Perm/Wash^TM^ Buffer (BD Biosciences). Finally, cells were washed with Perm/Wash^TM^ Buffer (BD Biosciences) and fixed with 1% paraformaldehyde (Sigma). Data was collected on an LSR II (BD Biosciences) and analyzed using FlowJo software (Version 9.7.6, Treestar Inc., Ashland, Oregon).

To evaluate T cell exhaustion, cryopreserved PBMCs and MLN lymphocytes were thawed and washed with R10, then stained with LIVE/DEAD® Aqua (ThermoFisher®). Cells were subsequently surface stained with an antibody cocktail consisting of CD3 BV 650 (Sp34-2, BD Biosciences), CD4 BV605 (OKT4, BD Biosciences), CD25 APC-R700 (2A3, BD), CD8 BV710 (RPA-T8, BD Biosciences), CD28 PE-CF594 (CD28.2, BD Biosciences), CD95 PerCP-eFluor710 (Dx2, ThermoFisher®), PD-1 BV785 (EHI12.2H7, BioLegend), TIGIT FITC (MBSA43, ThermoFisher®), SLAM BV421 (A12, BD), CTLA-4 PECy5 (BNI3, BD), and LAG-3 PE (polyclonal, R&D) in Brilliant Stain buffer (BD Biosciences). Fluorescence minus one (FMO) controls were included for each of the exhaustion markers, PD-1, TIGIT, SLAM, CTLA-4, and LAG-3, to more accurately determine their expression. Cells were subsequently washed with Brilliant Stain buffer (BD Biosciences) and fixed with 1% paraformaldehyde (Sigma). Finally, data was collected using an LSR II (BD Biosciences) and analyzed on FlowJo software (Version 9.7.6, Treestar Inc., Ashland, Oregon).

### Intracellular Cytokine Staining (ICS) of gut mucosa

Intraepithelial and lamina propria lymphocytes were isolated from colon biopsies and stimulated in the presence of brefeldin A (Sigma) and CD107a antibody (eBioH4A3; eBioscience), as previously described [40]. Cell were assessed for viability (Life Technologies) and stained using surface and intracellular/intranuclear markers as previously described [40]. All samples were acquired on an LSRII (BD Biosciences) and analyzed using FlowJo software version 9.9.4 (FlowJo; LLC). Gating schemes are described previously [40]. Briefly, CD4^+^ Tregs were designated by coexpression of CD25 and FoxP3 and Th17 cells were defiend by IL-17 production.

### Statistical Analyses

Statistical differences between multiple groups were calculated using a Dunn’s multiple comparisons test, while statistical comparisons between two groups were determined using a two-sided Mann-Whitney. Statistical differences in viral load, CD4^+^ T cell counts, or immune responses between timepoints was calculated using a Wilcoxon matched-pairs signed rank test. Viral burden was determined by calculating the area under the curve of each animal’s viral load graph. Correlations between immune responses and viral burden were determined by a Spearman’s rank correlation test. When necessary, P values were adjusted for multiple comparisons using the Benjamini-Hochberg method. A P value of ≤ 0.05 was considered significant for each test. All calculations were performed using GraphPad Prism software (Version 7, GraphPad Software, San Diego, CA).

## Acknowledgements

The authors would like to thank all the veterinary and research support staff of the Washington National Primate Research Center (WaNPRC), with special thanks given to Solomon Wangari, Drew May, Dr. Jennifer Lane, Dr. Cassie Moats, Dr. Jeremy Smedley, and Dr. Robert Murnane. We also wish to thank Thomas Lewis and Dr. Patience Murapa for their early work in determining the optimal adjuvant combination for use in this study. Gag and Env peptides and SIV gp130 proteins were generously provided by the NIH AIDS Research and Reference Reagent Program.

---

S1 Fig. Animals in each group demonstrate similar plasma viral loads, CD4 T cell counts, and ART responsiveness.

**(A)** Plasma viral loads as determined by RT-q-PCR for the mock (black circles), MAG + LT (red squares) and MAG + AC (purple triangles) groups, shown are medians and interquartile ranges. The dashed line indicates the assay limit of detection (30 viral RNA copies/1mL of plasma) and the dotted line indicates the threshold for control of virus replication**. (B)** Percent of baseline CD4 T cell counts were calculated for the mock, MAG + LT and MAG + AC groups over time by dividing the absolute CD4 count at a timepoint by the absolute CD4 count at 0 wpi and multiplying by 100. Shown are medians and interquartile ranges. The dotted line indicates 50% of baseline CD4 T cells. CD4 T cell counts were obtained using a Beckman Coulter® AC*T^TM^ 5diff hematology analyzer. **(C)** Shown is the decrease of each animals’ viral loads between pre-ART (6 wpi) and pre-vaccination (32 wpi). Statistical analyses were performed using a Wilcoxon matched-pairs signed rank test; results are considered significant if P ≤ 0.05. **(D)** Shown is the restoration of each animals’ percent of baseline CD4 T cell counts between pre-ART (6 wpi) and pre-vaccination (32 wpi). Statistical analyses were performed using a Wilcoxon matched-pairs signed rank test; results are considered significant if P ≤ 0.05.

S2 Fig. No differences were observed between groups in Gag-specific CD4^+^ T cell responses in PBMC and MLN.

**(A)** PBMCs were thawed and stimulated with Gag peptides, and expression of IL-2, IFNγ, TNFα and CD107a/GzB were quantified using intracellular cytokine staining. Shown are the medians and interquartile ranges of each group’s SIV-Gag specific T cell response. **(B)** Lymphocytes isolated from MLN were thawed and stimulated with Gag peptides, and expression of IL-2, IFNγ, TNFα and CD107a/GzB were quantified using intracellular cytokine staining. Shown are the medians and interquartile ranges of each group’s SIV-Gag specific T cell response. **(A, B) Statistics**. Statistical comparisons between baseline and post-vaccination timepoints within a group were calculated using a Wilcoxon matched-pairs signed rank test. A Dunn’s multiple comparisons test was used when making multiple comparisons between vaccine groups and the mock group. Results are considered significant if P ≤ 0.05.

S3 Fig. No differences were observed among groups in Env-specific CD8^+^ or CD4^+^ T cell responses in PBMC.

**(A-B)** PBMCs were thawed and stimulated with Env peptides, and expression of IL-2, IFNγ, TNFα and CD107a/GzB were quantified using intracellular cytokine staining. Shown are the medians and interquartile ranges of each group’s SIV-Env specific T cell response. **(A, B) Statistics**. Statistical comparisons between baseline and post-vaccination timepoints within a group were calculated using a Wilcoxon matched-pairs signed rank test. A Dunn’s multiple comparisons test was used when making multiple comparisons between vaccine groups and the mock group. Results are considered significant if P ≤ 0.05.

S4 Fig. No differences were observed among groups in Env-specific CD8^+^ or CD4^+^ T cell responses in MLN.

**(A-B)** Lymphocytes isolated from MLNs were thawed and stimulated with Env peptides, and expression of IL-2, IFNγ, TNFα and CD107a/GzB were quantified using intracellular cytokine staining. Shown are the medians and interquartile ranges of each group’s SIV-Env specific T cell response. **(A, B) Statistics**. Statistical comparisons between baseline and post-vaccination timepoints within a group were calculated using a Wilcoxon matched-pairs signed rank test. A Dunn’s multiple comparisons test was used when making multiple comparisons between vaccine groups and the mock group. Results are considered significant if P ≤ 0.05.

S5 Fig. CD4^+^ T cell counts corresponded with virus burden in plasma.

**(A-C)** Shown are the percent of baseline CD4^+^ T cell counts for each individual animal in the mock, MAG + LT and MAG + AC groups over time. Percent of baseline CD4^+^ T cell counts were calculated for the mock, MAG + LT and MAG + AC groups over time by dividing the absolute CD4^+^ count at a timepoint by the absolute CD4^+^ count at 0 wpi and multiplying by 100. The dotted line indicates 50% of baseline CD4^+^ T cells. CD4^+^ T cell counts were obtained using a Beckman Coulter® AC*T^TM^ 5diff hematology analyzer. **(D)** Graphed are the median and interquartile range of controllers’ and noncontrollers’ percent of baseline CD4^+^ counts.

S6 Fig. No difference between controllers/noncontrollers in Gag-specific IL-2^+^ and CD107a^+^GzB^+^ CD8^+^ T cells in PBMC or Gag-specific IFNγ^+^, TNFα^+^, and CD107a^+^GzB^+^ CD8^+^ T cells in MLN.

**(A-E)** PBMCs and MLNs were thawed and stimulated with Gag peptides, and expression of cytokines was quantified using intracellular cytokine staining. Shown are the medians and interquartile ranges of the SIV Gag-specific CD8^+^ T cell responses of controllers and noncontrollers, with individual responses layered over each bar at a post-vaccination timepoint (50 wpi) and during ATI (62 wpi for PBMC and 66 wpi for MLN). Statistical differences between controllers and noncontrollers at each timepoint were assessed using a Mann Whitney t test and the Benjamini-Hochberg method was used to adjust P values. Results are considered significant if P ≤ 0.05.

S7 Fig. Controllers and noncontrollers demonstrate similar levels of SIV Gag-specific CD4^+^ T cell responses in the PBMC and MLN post-vaccination and during ATI.

**(A-B)** PBMCs and MLNs were thawed and stimulated with Gag peptides, and expression of cytokines was quantified using intracellular cytokine staining. Shown are the medians and interquartile ranges of the SIV Gag-specific CD4^+^ T cell responses of controllers and noncontrollers, with individual responses layered over each bar at a post-vaccination timepoint (50 wpi) and during ATI (62 wpi for PBMC and 66 wpi for MLN). Statistical differences between controllers and noncontrollers at each timepoint were assessed using a Mann Whitney t test and the Benjamini-Hochberg method was used to adjust P values. Results are considered significant if P ≤ 0.05.

S8 Fig. Noncontrollers exhibited a trend towards higher titers of Env-specific IgG compared to controllers.

The magnitude of the SIV Env-specific IgG response in the plasma was measured by ELISA, using SIV gp130 as the capture antigen. Shown are medians and interquartile ranges.

S9 Fig. Baseline frequencies of Th17 and Treg cells may play a role in determining therapeutic outcome during ATI.

**(A-B)** Lymphocytes were isolated from colon biopsies and expression of Th17 and Treg markers was quantified using intracellular cytokine staining on fresh cells. A Spearman rank correlation test was used to determine P and r values. Shown are the Benjamini-Hochberg adjusted P values, results are considered significant if P ≤ 0.05.

